# Fly wing evolutionary rate is a near-isometric function of mutational variation

**DOI:** 10.1101/2020.08.27.268938

**Authors:** David Houle, Geir H. Bolstad, Thomas F. Hansen

## Abstract

If there is abundant mutational and standing genetic variation, most expect that the rate of evolution would be driven primarily by natural selection, and potentially be independent of current variability or variation. Contrary to this expectation, we (H17: Houle et al. 2017 Nature 548:447) found surprisingly strong scaling relationships with slopes near one between mutational variance, standing genetic variance and macro-evolutionary rate in Drosophilid wing traits. Jiang and Zhang (J&Z20: 2020 Evolution https://doi.org/10.1111/evo.14076) have challenged these results and our interpretation of them. J&Z20 showed that the method used in H17 to estimate the scaling relationship between variation at different biological levels is uninformative. Using an alternative method, they estimated that the scaling relationship has a slope substantially less than one, and propose a variant of our neutral subset hypothesis to explain this. Here we use simulations to confirm J&Z20’s finding that the H17 method for estimating scaling of variances is uninformative. The simulations also show their alternative method for estimating scaling is likely to be seriously biased towards lower scaling relationships. We propose and verify an alternative approach to calculating scaling relationships based on independently estimated variance matrices, which we call the Q method. Simulations and reanalyses of the Drosophilid data set using the Q method suggests that our original estimates of the scaling relationship were close to the true value. We propose an analytical version of the neutral subset model, and show that it can indeed explain any scaling slope by varying assumptions about the pattern of pleiotropy. We continue to regard neutral subset models as implausible for wing shape in Drosophilids due to the likelihood that wing shape is subject to direct selection. Hybrid models in which pleiotropy reduces the available genetic and mutational variation, and a combination of selection and drift controls the change in species means seem more biologically promising.

## Introduction

High levels of standing genetic variance and mutational input predict that quantitative traits should not be evolutionarily constrained on macro-evolutionary time scales. Given this, we expect that the rate of evolution in traits subject to natural selection would be driven primarily by selection and potentially be independent of current variability or variation. Contrary to this expectation, we recently found surprisingly strong scaling relationships between mutational variance, standing genetic variance and macro-evolutionary rate in Drosophila wing traits (Houle et al. 2017, hereafter H17). This result, if correct, challenges current theory.

Jiang and Zhang (2020, hereafter J&Z) have challenged some aspects of the results and interpretation in H17.. J&Z20 make four main points. First, they show that the method that we adopted to estimate the scaling relationship between variation at different biological levels is uninformative and biased. Second, they claim that we have overestimated the slope of the relationship between mutation and the rate of evolution. Third they show that a specific model that invokes mutational pleiotropy can explain a non-isometric scaling between mutational variance and the rate of evolution. Fourth, they suggest that this mutational model is more plausible than a similar model that we termed implausible. We take up each of these points in turn.

In this contribution, we consider these challenges to our results using a combination of additional analyses of results in H17 and simulation. In addition, we offer an analytical version of the model that J&Z20 propose to explain non-isometric scaling. We confirm that J&Z20 are correct that our method for estimating scaling of variances is uninformative. The alternative method proposed by J&Z20, however, is likely to produce downwardly biased slope estimates. We propose an alternative method for calculating the scaling relationship which is more accurate than either our original method or that of J&Z20. The new method suggests that our original estimates of the scaling relationship near one are close to the true value.

### Background to the controversy

In H17 we studied the relationship between mutational variation, standing genetic variance, and the rate of evolution among species for a suite of wing shape and size traits in the family Drosophilidae. Mutational and additive genetic variation were estimated in one species, *Drosophila melanogaster*, and summarized in an **M** and a **G** matrix respectively. In addition we estimated the rate of divergence in wing shape among 110 species in the family Drosophilidae, summarized in a rate matrix, **R**. Each matrix summarized variation or evolution for a total of 21 traits, log(centroid size) and 20 wing shape variables. Consequently, the resulting variance matrices have high dimensionality, and can be similar and different from each other in a multitude of ways. When we compared variation in a set of traits that spanned the phenotype space, such as in Figure 4 of H17, both similarities and differences seemed evident.

Principal components analysis (PCA) is a natural way to choose traits in shape space, as it identifies statistically independent shape axes that form a basis for shape space. However PCA of any combination of our data would generate a valid basis. A priori, we know that the choice of basis potentially affects our results. PCA maximizes the variance in eigenvalues, the variances along each linear combination of measurements used to define a traits. If we have two variance matrices **A** and **B**, and we would like to predict variation in **B** from **A**, using eigenvalue decomposition of **A** to define traits will maximize the variance among predictor variables, while using **B** to define traits will maximize the variance among dependent variables. Variance matrices are always estimated with error, so the maximization entailed in PCA will also incorporate error variance into the most extreme eigenvalues, biasing the variance and range of eigenvalues relative to the true values (Hill and Thompson 1978; Johnstone 2001). In the context of a linear regression, maximizing the variation in the predictor variable will bias the estimated slope downwards, while maximizing the variation in the dependent variable will bias the estimated slope downwards (Hayashi et al. 2018). Examples of these biases are presented in Figure S1

In H17, we treat the mutational variability summarized in **M** as a potential cause of the variation in evolutionary rate summarized in **R**. Thus choosing traits based on PCA of **M** will bias the estimated slope downwards. Initial analyses using **M** and **R** confirmed the direction of these biases; defining traits based on **M** resulted in substantially lower estimates of slope than definition of traits based on **R**. We (H17) therefore chose traits as the eigenvectors of a matrix 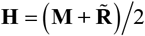, where 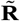 is **R** scaled to the same size as **M**. We refer to choosing the basis for comparison using the predictor matrix as the M method, and using the scaled average of the predictor and dependent matrices as the H method. We knew from previous work on sampling error of variance matrices that the M method creates a bias (Hayashi et al. 2018), and naively assumed that the H method would reduce or eliminate that bias.

## Methods

### Analyses of empirical data

In this contribution we utilize the wing shape variation in four different empirical variance matrix estimates drawn from the data in Houle et al. (2017; H17). We consider only the mutation matrix generated from homozygous line means, **M**_hom_ here, which we will refer to as **M**. The matrices **G** and **R** were presented in H17. We obtained 1,000 matrix estimates from the sampling distribution of **M**, **G** and **R** using the REML-MVN approach (Meyer and Houle 2013; Houle and Meyer 2015). The final matrix is the pooled phenotypic variance matrix **P**. To generate this matrix, we calculated the variance matrix among species- and sex-standardized wing shape variables in the 110 Drosphilid species used to calculate **R**. We calculated 1,000 such estimates of **P** from bootstrap samples at the level of species.

**M** is not full rank, and has 18 significant dimensions. We focus on just the first *k*=17 dimensions of the **M** phenotype space, as the corresponding eigenvalues of **M** approximate an exponential distribution. To compare variation among matrices, we first projected each matrix (for example **A**) into the 17-dimensional space spanned by **M** using

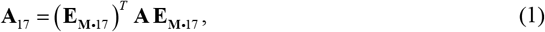

where **E_M•17_** contains the first 17 eigenvectors of **M**. These eigenvectors form a basis for the 17 dimensional phenotype space. We then scaled each matrix to unit size using **Ã** = **A**_17_/tr(**A**_17_), where tr() indicates the trace of the matrix. 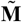 is a diagonal matrix containing the first 17 scaled eigenvalues of **M**.

We investigated five possible phenotypic bases to investigate the relationship between 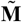 and 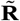, the eigenvectors of the matrices **M**, **R**, **G**, **P**, and **H**., For example, to use a basis estimated from **A**, eigendecompose **A** to yield the eigenvectors **E_A_**, rotate 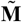 and 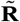 into that space using 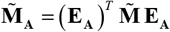 and 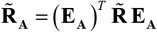. The final step is to regress the diagonal elements of **R_A_** on the diagonal elements of **M_A_** on a log_10_ scale. The sampling variation of the regression slopes were estimated from the distribution of estimated slopes over the 1,000 replicate estimates of 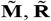, and **A**.

In H17, we did not directly estimate the scaling relationships by simple regression. Instead we fit a phylogenetic mixed model that better incorporated the structure of the data. Additional details are given in H17. For comparison with the direct regression estimates, we repeated these analyses for each of the five bases in this manuscript.

Distances between variance matrices were calculated using the geodesic distance

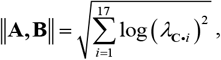

where **A** and **B** are the matrices to compare, *λ*_**C**•*i*_ is the *i*th ranked eigenvalue of **C** = (**A**)^−1^ **B** (Mitteroecker and Bookstein 2009).

Statistical analyses were carried out in SAS (SAS Institute 2016), with the exception of estimation of the phylogenetic mixed model estimates of variance scaling relationships, which were fitted using the Template Model Builder (Kristensen et al. 2015) implemented in R (R Core Team 2020).

### Simulations

Our simulations draw sample matrices from a Wishart distribution, with a parameter matrix **Â**, denoted with a carat. To generate the simulated parameter matrices including 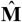 and 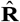 we first generated diagonal matrices with log_10_ eigenvalues

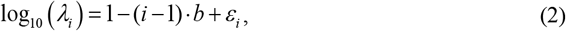

where *λ* is eigenvalue, *i* is eigenvalue rank, *b* is the rate of decrease in eigenvalues with rank, and 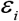 is a random variate. The eigenvalue error, 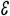 was drawn from a normal distribution with zero mean, and variance *v*. To match the observed data, we generated mutation parameter matrices, 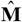 using *b*=-0.154 and *v*=0.007.

To generate the dependent parameter matrices, 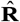 and independent basis matrices 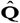, with differing relationships to 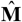 we randomly rotated 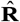 and 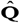 relative to 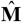. We first generate a diagonal matrix **X** using equation (1), which is in expectation maximally similar to 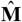 subject to the choice of *b* and *v*. This initial matrix is then perturbed using a series of *t* Givens rotations (Kirkpatrick and Meyer 2004) of angle *θ* in radians around randomly chosen planes in phenotype space. To perform the *i*th such rotation, we first generated two random vectors in phenotype space by drawing *k*= 17 variates from a standard normal distribution 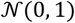. Gram-Schmidt orthonormalization of these two vectors gives two orthogonal basis vectors which were formed into a 17 x 2 matrix **W**_i_. The *i*th rotation matrix is

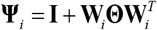

where **I** is the 17 x 17 identity matrix, ^*T*^ indicates transpose and

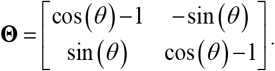

The final rotated parameter matrix is

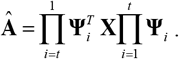

The similarity of 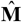 to the other parameter matrices is jointly determined by *ε, θ* and *t*. For all simulations we chose *t*=17, corresponding to the dimensionality of the matrices.

The true regression slope of variation in 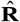 on 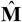 was calculated using the eigenvectors of 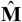 to define traits. The estimated slope is a non-linear function of the parameters, so for presentation we label samples according to the true mean slope generated.

The final step is to generate sample estimates **M**_*j*_ **R**_*j*_ and **Q**_*j*_ from a Wishart distribution. The Wishart parameters included the appropriate parameter matrix and the degrees of freedom for the sample matrices. Degrees of freedom were varied as described in the Results section.

## Results

### Simulations of the H method

We first compared the H method for choosing traits used in H17, and the M method advocated in J&Z20 using a different simulation approach from that used in J&Z20. As shown in Figure S2, we confirm J&Z20’s finding that the H method is largely insensitive to the true scaling relationship, always recovering a slope near 1. The M method is a valid estimator of relationship, although it is biased downwards, as expected.

### Reanalysis of data from H17

To provide the full context for the analysis of the H17 data set, we reanalyzed several aspects that data set. Houle et al. (2017; abbreviated H17) estimated the scaling relationship between log_10_ variation in **R** as a function of log_10_ variation in **M** using a phylogenetic mixed model, while (Jiang and Zhang 2020; abbreviated J&Z20) applied a simple regression-based approach. We recalculated the scaling relationship using the least squares and the mixed model approaches using the eigenvectors of each of the available matrices as a basis, with the results shown in the first few columns of Table 1. Differences between simple regression and phylogenetic mixed models are small relative to the variation generated by choice of basis. The difference between the two approaches statistical significance only when **R** is used to estimate the basis vectors. In the remainder of this paper, we focus on investigating how the choice of basis influences the estimate of scaling calculated by simple regression, as J&Z20 did.

**Table 1.**
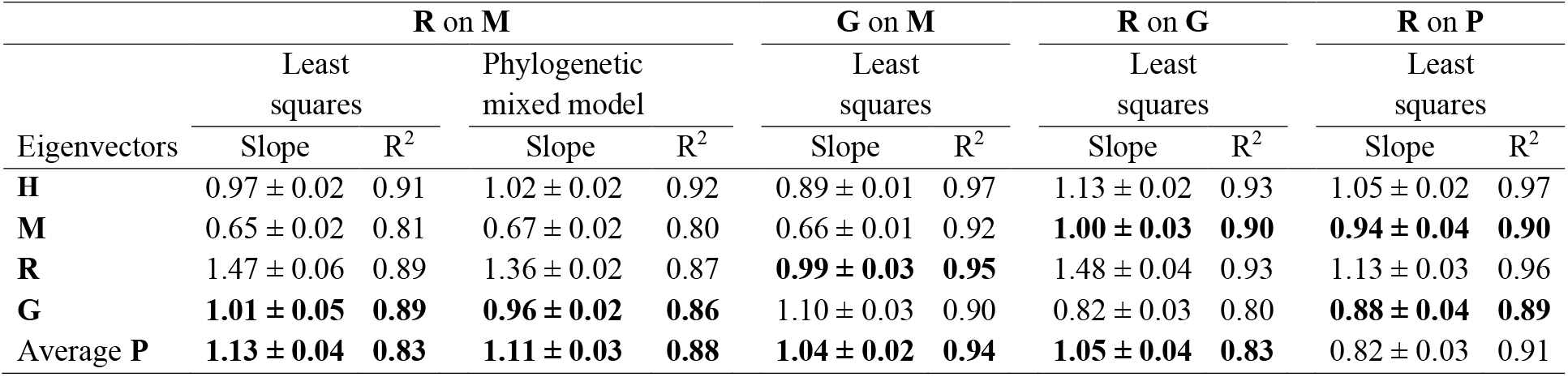
Regression of log_10_ **R** variation on log_10_ **M** variation using the first 17 eigenvectors of five different sample matrices as the basis for comparisons. Bold-faced estimates are not biased by sampling variation within matrices. Slopes were estimated using least squares regression or a phylogenetic mixed model (Houle et al. 2017).

| Eigenvectors | **R on M** |  |  |  | **G on M** |  | **R on G** |  | **R on P** |  |
| --- | --- | --- | --- | --- | --- | --- | --- | --- | --- | --- |
|  | Least squares |  | Phylogenetic mixed model |  | Least squares |  | Least squares |  | Least squares |  |
|  | Slope | R<sup>2</sup> | Slope | R<sup>2</sup> | Slope | R<sup>2</sup> | Slope | R<sup>2</sup> | Slope | R<sup>2</sup> |
| **H** | 0.97 ± 0.02 | 0.91 | 1.02 ± 0.02 | 0.92 | 0.89 ± 0.01 | 0.97 | 1.13 ± 0.02 | 0.93 | 1.05 ± 0.02 | 0.97 |
| **M** | 0.65 ± 0.02 | 0.81 | 0.67 ± 0.02 | 0.80 | 0.66 ± 0.01 | 0.92 | **1.00 ± 0.03** | **0.90** | **0.94 ± 0.04** | **0.90** |
| **R** | 1.47 ± 0.06 | 0.89 | 1.36 ± 0.02 | 0.87 | **0.99 ± 0.03** | **0.95** | 1.48 ± 0.04 | 0.93 | 1.13 ± 0.03 | 0.96 |
| **G** | **1.01 ± 0.05** | **0.89** | **0.96 ± 0.02** | **0.86** | 1.10 ± 0.03 | 0.90 | 0.82 ± 0.03 | 0.80 | **0.88 ± 0.04** | **0.89** |
| Average **P** | **1.13 ± 0.04** | **0.83** | **1.11 ± 0.03** | **0.88** | **1.04 ± 0.02** | **0.94** | **1.05 ± 0.04** | **0.83** | 0.82 ± 0.03 | 0.91 |

The rightmost columns in Table 1 show the least-squares regressions for each combination of a lower level matrix as a predictor of **G** and **R**. The H method takes the scaled average of the dependent and predictor matrices. For each scaling relationship, using the predictor matrix to choose the basis results in the lowest estimated slope, while choosing the basis using the dependent matrix gives the highest estimate of slope. The estimates using an independently estimated basis, for example, the relationship between **M** and **G** using a basis estimated from **P**, always give slopes that are more similar to each other than they are to the slopes estimated along bases from the dependent or independent matrices. This suggests that the realized biases from the sampling variation in the dependent or independent matrices have a substantial effect on the results. This in turn suggests that perhaps estimates based on a basis estimated from an independent matrix may be more accurate. We call this approach the Q method, where Q can be any independently-estimated matrix. Below, we describe simulations to test whether this is true.

High-dimensional variance matrices can differ in a very wide number of aspects. Previous analyses (Mezey and Houle 2005; Houle and Fierst 2013; Houle et al. 2017) revealed that the eigenvalues of **M**, **G** and **R** were well-approximated by an exponential distribution. We tested whether this reflected a common relationship between eigenvalue and eigenvector rank in our four sample matrices. We first estimated the first 17 eigenvalues of the **M**, **G**, **R** and **P** matrices in separate principal component analyses (PCAs). We then performed an analysis of covariance of these eigenvalues with matrix as a fixed effect and eigenvalue rank as a continuous predictor, with results shown in Table 2. As expected from Figure 4 in H17, the effects of eigenvector rank and matrix are highly significant, but there is no significant difference in slope between matrices (P=0.15). Overall, the model explains 99% of the variance in eigenvalues. The most divergent slope is that for **G**, and removing **G** from the analysis increases the rank by matrix interaction *P* value to 0.58. The estimated slope of eigenvalue on rank for **G** is −0.139 ± 0.0040 (S.E.), while the pooled slope for the other three matrices is −0.156 ± 0.0048. The residuals from the common slope were heteroscedastic, reflecting some departure from the exponential distribution for **R** and **P**. This is reflected in the residual variance from separate regression models, with 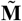 and 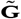 having residual variances of 0.007, while residual variance for 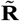 was 0.027, and that for 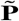 was 0.054.

We also characterized the geodesic distances between our size-standardized variance matrices, with results shown in Figure 1. Once standardized to the same size, all of the variance matrices are about equally distant from each other across the manifold of possible variance matrices.

**Figure 1.**
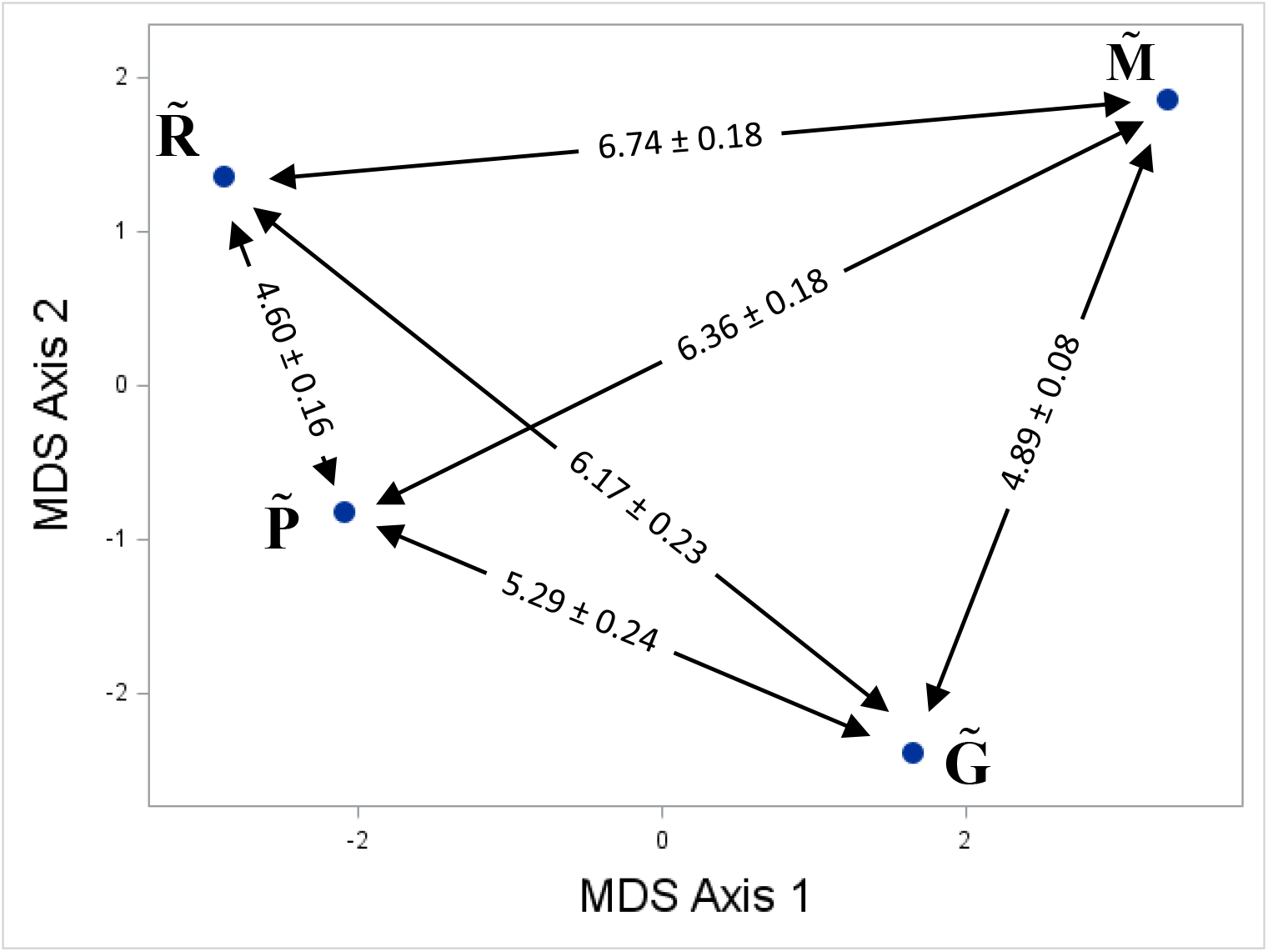
Position of 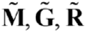 and 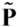 in on the manifold of possible variance matrices. Positions projected into two-dimensional space using non-metric multidimensional scaling. Positions in two dimension explains 91.1% of the distance relationships among matrices. Geodesic distances calculated from matrices standardized to size 1 ± standard error.

### Rationale for the Q method

Our goal is to choose a basis, a set of orthogonal traits that defines the multi-dimensional data space, that enables accurate estimates of regression slope between variation in two sample variance matrices. Based on the results shown in Table 1, we hypothesized that the use of an independent matrix to choose the basis vectors might result in more accurate estimates of the scaling relationship between a predictor and dependent matrix than a choice dependent on the predictor and dependent matrices. We refer to this as the Q method, where **Q** can represent any variance matrix of the same or larger dimension as the matrices whose variance are compared. J&Z20 advocate adopting the M method in which the predictor matrix is used to determine the set of traits on which variation is compared. The M method is known a priori to generate biased estimates, and our simulations and those in J&Z20 confirm the existence of the bias. J&Z20 show that the sampling variation in the estimate of **M** that H17 used (Houle and Fierst 2013) is too small to give rise to a substantial bias. However, there are other potentially more important sources of variation. Under the hypothesis that mutational variance shapes the rate of evolution, the average mutational variation over the entire clade, call it 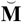, is the relevant parameter matrix, rather than the **M** matrix within one particular population and species. Biological sampling of the genotypes used to estimate **M**_i_ and evolution of the **M** matrix within the clade likely contribute more to discrepancies between **M**_i_ and 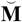 than sampling variation.

There is direct evidence for this biological sampling. The **M**_i_ matrix used in H17 (Houle and Fierst 2013) is the average of estimates from two different inbred genotypes derived from one natural population of *Drosophila melanogaster*. These two genotypes differed significantly in mutation at both the genomic (Haag-Liautard et al. 2007) and the phenotypic level, with striking differences between the two estimates in many directions in phenotype space (Houle and Fierst 2013). For example, of the 18 dimensions with significant mutational variance in the pooled sample, only 10 dimensions had significant genetic variance in both matrices. Even in those dimensions with significant variation in both matrices, the amounts of variance differed by a factor of as much as 108. Thus, there is clear evidence for variation in **M** within one population of *D. melanogaster* from this one experiment. It is thus very likely that 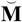 may be different from the single estimate of **M**_i_ used in H17. On the other hand, this expected discrepancy between **M**_i_ and 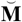 is one reason why the strong scaling relationships observed in H17 are unexpected. If the discrepancies are not large, then the M method might perform well, despite some degree of bias.

In the H17 study, **G** and **P** are estimated independently of **M**, and could be used in the Q method. If **M** has a similar relationship to **G** and **P** as it does to **R**, as suggested by the overall similarities of the matrices in H17, **G** and **P** are potentially better representations of the eigenvectors of 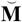 than the eigenvectors of **M**. **G** potentially integrates the mutational variation across genotypes within the sampled population. The pooled **P** across species potentially integrates mutational variation across all species used to estimate **R**. On the other hand, if a **Q** matrix does not have a strong relationship to the predictor matrix, this could result in other biases and inaccuracies.

### Simulations of the Q method

We carried out additional simulations to evaluate the bias and precision of the Q and M methods under different assumptions about the true scaling relationship and the accuracy with which matrix is estimated. For simplicity, we refer to predictor matrices as **M** and dependent matrices as **R**, although our results apply equally well to any combination of predictor and dependent matrices.

We judged that the eigenvalue distributions in our estimated matrices were similar enough that we could assume a common regression slope in all parameter matrices in our simulations without affecting the applicability of conclusions to the H17 data. The slope of eigenvalue on rank in each simulated parameter matrix was set to −0.16, plus a normally distributed deviation with mean 0 and variance 0.007 for 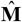, and 0.027 for 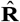. Initially we used 160 for the degrees of freedom when drawing sample matrices from the Wishart distribution.

The upper limit on the degrees of freedom with which to draw sample matrices from the Wishart distribution is set by the sampling variance of each matrix. Therefore, we determined the relationship between residuals of the regression of log_10_ eigenvalues on eigenvector rank and the degrees of freedom (df) parameter in the Wishart distribution with results shown in Figure S3. The regression of residual variance on df is log_10_ (sampling variance) = −0.553 – 0.293(log_2_ (df)) and has *R*^2^ = 0.99. The average sampling variance of eigenvalues for each of the estimated matrices in H17, standardized to unit size, is shown in Table S1 along with the predicted df that would result in that average sampling variance. **M** is the most precisely estimated matrix, with an equivalent df=1246, while **R** is the least precisely estimated, with equivalent df=101. The equivalent df for the phenotypic matrix **P** of 142 is rather low due to the sampling of species from which **P** was been estimated. Within-species sampling variation of **P** is quite small, as the average sample size per species is 192 individuals. **P** incorporates among-species biological variation that has not been estimated for **G** or **M**.

Figure 2 illustrates the effects of sampling from the parameter matrices, 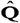 and 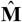, on the estimated scaling relationship between **R**_*i*_ and **M**_*i*_ using the M and Q methods. To vary scaling relationships, we generated a statistically independent parameter matrix, 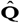, with the same algorithm we used to generate 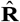 and then randomly rotated those estimates relative to **R** and **M**. When **M**_*i*_ differs from 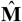 only due to the empirically-calibrated level of sampling variation (Wishart df=1280, Fig. 3, upper panels), the M method gives very accurate estimates of the true scaling relationship, consistent with the results of J&Z20. However, when there is substantial biological variation on top of sampling variation (Wishart df=30, Fig. 2, lower panels), the M method is downwardly biased by a substantial amount. The Q method tends to have a slight positive bias, with the exception of the case when 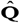 and 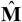 are unrelated. The sampling variation in **Q**_*i*_ has a much more modest effect on the scaling relationship than does sampling of **M**_*i*_. There is an interaction between sampling variation in **M**_*i*_ and **Q**_*i*_. Comparison of the top panels with the lower panels shows that at both levels of sampling of **Q**_*i*_, the Q method bias is larger when **M**_*i*_ is closer to 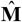 than when it is distant due to sampling.

**Figure 2.**
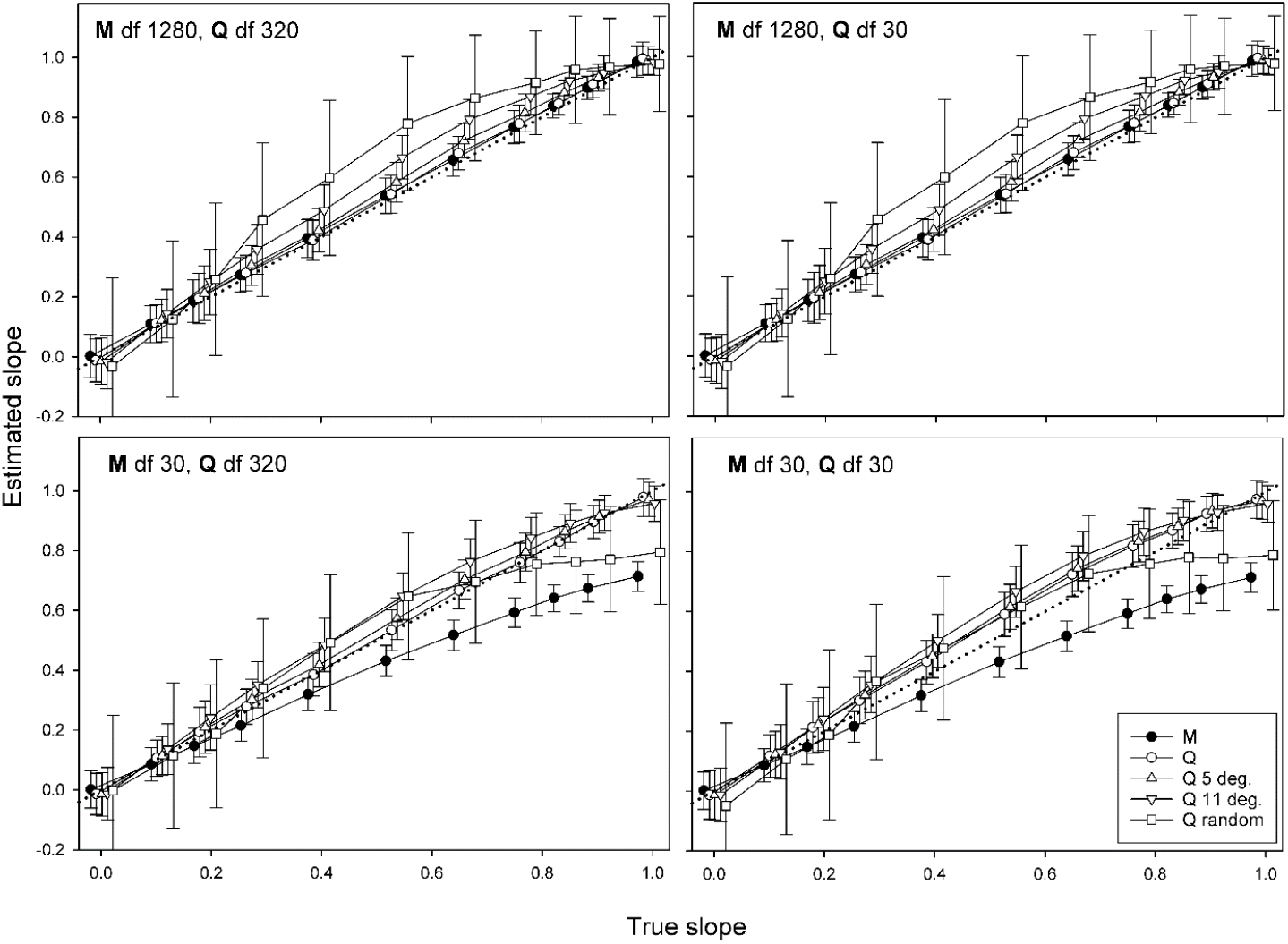
Estimated scaling relationships between **R**_*i*_ and **M**_*i*_, ± standard error, when the M and Q methods are used to define a basis. True slopes are constant across basis assignment methods, symbols are offset to improve visibility. Dotted line is the 1:1 line. M = M method; Q = Q method when 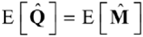; Q 5 deg. = as in Q, but 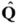 subjected to random rotations by 4.5 degrees; Q 11 deg. = as in Q, but 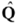 subjected to random rotations by 10.8 degrees; Q random = orientation of 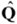 random with respect to 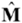.

Clearly, the best method for choosing a basis depends on the details of the relationship between the parameter and sample matrices, as shown by the contrast between the performance of the M method between the rows of Figure 2, and between the sample matrices to each other, as shown by the dependence of the performance of the Q method on the sampling variation of **M**_*i*_.

The results of the comparisons between **M**, **R**, **G** and **P** from the H17 data sets provide the opportunity to choose a set of simulation parameters that approximate these relationships. The R method gives a very high slope. The M method gives a markedly lower estimate, while the two Q method estimates, based on **G** and **P**, give an intermediate slope. These results are consistent with a distant relationship between **M**_*i*_ to 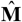 due to biological sampling, as suggested by the results in Figure 2. A second criterion is the proportion of variance explained by the scaling relationship. Figure S4 shows *R*^2^ values as a function of the relationship between 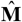 and 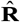, on the x-axis, for different choices of basis and sampling effort. *R*^2^ values greater than 0.8, consistent with the results in Table 1, can only be achieved when the true scaling relationship is greater than about 0.75 regardless of error in the estimated matrices. Finally, Figure 3 explores the geodesic distances expected between **M**_*i*_ and **Q**_*i*_ under various levels of sampling error. When **Q**_*i*_ is drawn from a random 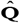, the geodesic distances are always much higher than the distances between the observed standardized matrices from H17 shown in Figure 1. When **Q**_*i*_ is drawn from 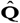 with closer relationships to 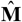, geodesic distances can fall with the observed range of estimates either when 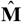 is well estimated and 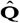 is rotated to be less similar to 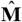, or when 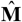 is inaccurately estimated and 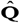 is more similar to 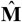. Taken together, these results suggest that the true scaling relationship is greater than 0.75, the biological sampling variance in **M**_*i*_ is large, while the sampling variances in potential **Q** matrices **G** and **P** are intermediate. 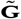 and 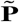 are both more similar to 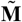 and 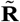 than 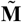 is to 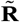.

**Figure 3.**
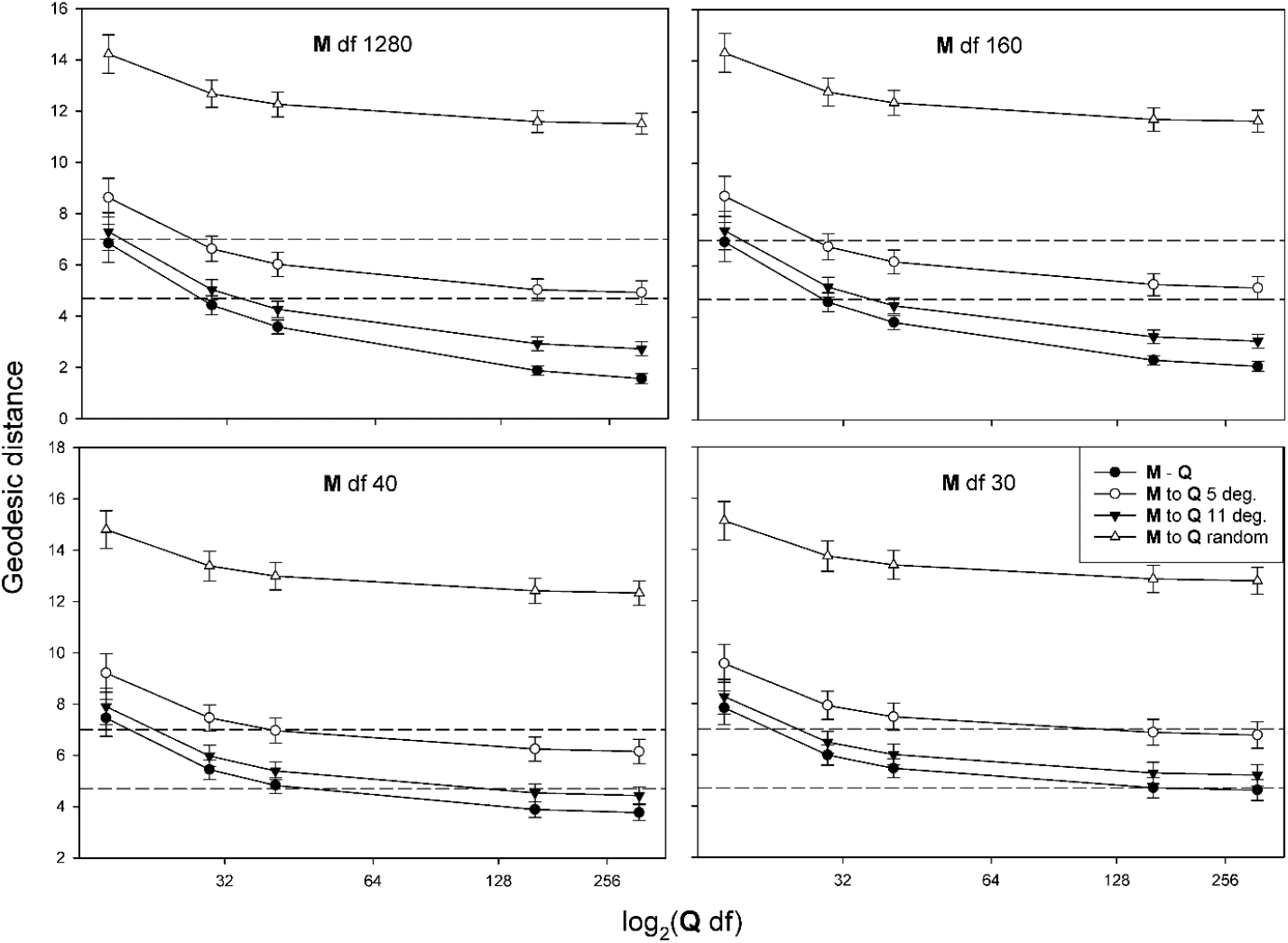
Geodesic distances between **M**_*i*_ and **Q**_*i*_ under various combinations of sampling error. Dashed reference lines show the range of distances among the observed matrices 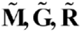 and 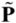 estimated from the data in H17. Legend as in Fig. 2.

Figures 4 and 5 show the bias and mean-squared error for various combinations of sampling degrees of freedom and relationships between **M**_*i*_ and **Q**_*i*_. In both figures, the upper left panel shows results when all of the deviation of **M**_*i*_ from 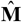 and **Q**_*i*_ from 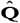 is due to sampling variance similar to the matrices in H17. Panels in the right column add a large component of biological variation to estimates of **Q**_*i*_. The middle and lower rows reflect the addition of a substantial component of biological variation to **M**_*i*_, consistent with the large discrepancy in estimated slope when using **M** versus **G** or **P** to choose the trait basis. Comparison of the left and right sides of the Figures shows that biological deviations of **Q**_*i*_ increase bias in the estimates a bit, but have essentially no effect on root mean squared error under the Q method. The addition of biological error to **M**_*i*_, however, has a large effect on the bias and root mean-square error of the M estimates. For the large values of the true slope consistent with high *R*^2^, realistic biological deviations of **M**_*i*_ from 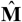 always raises the bias under the M method to levels higher than under the Q method. We cannot be sure which combinations of sampling df are most appropriate for the data in H17, but the simulated data sets consistent with all the empirical results suggest that the Q method will perform better than the M method for the matrices estimated in H17.

**Figure 4.**
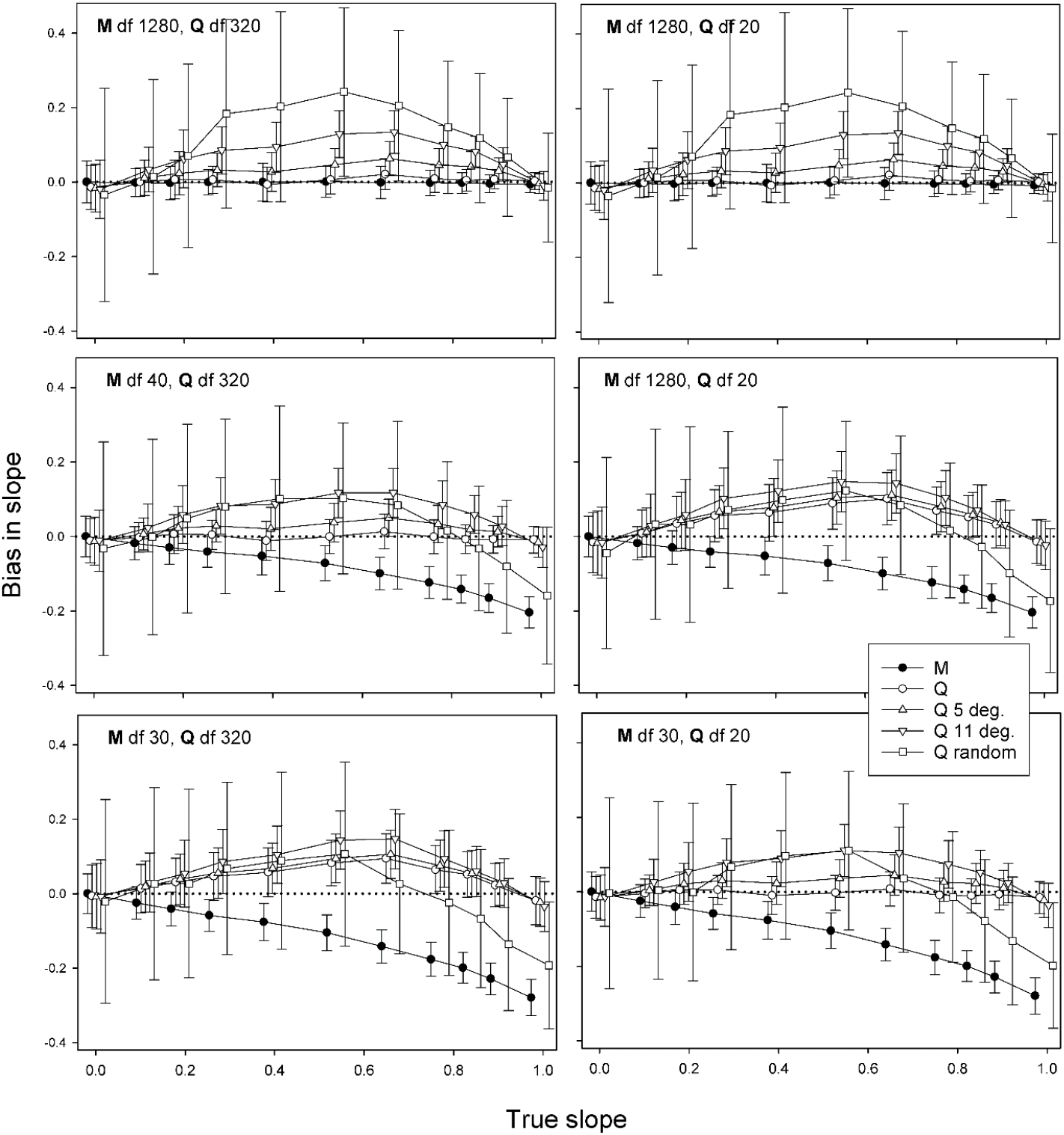
Bias in estimated scaling relationship, ± standard error, as a function of true relationship, sampling variation, and choice of basis method.

**Figure 5.**
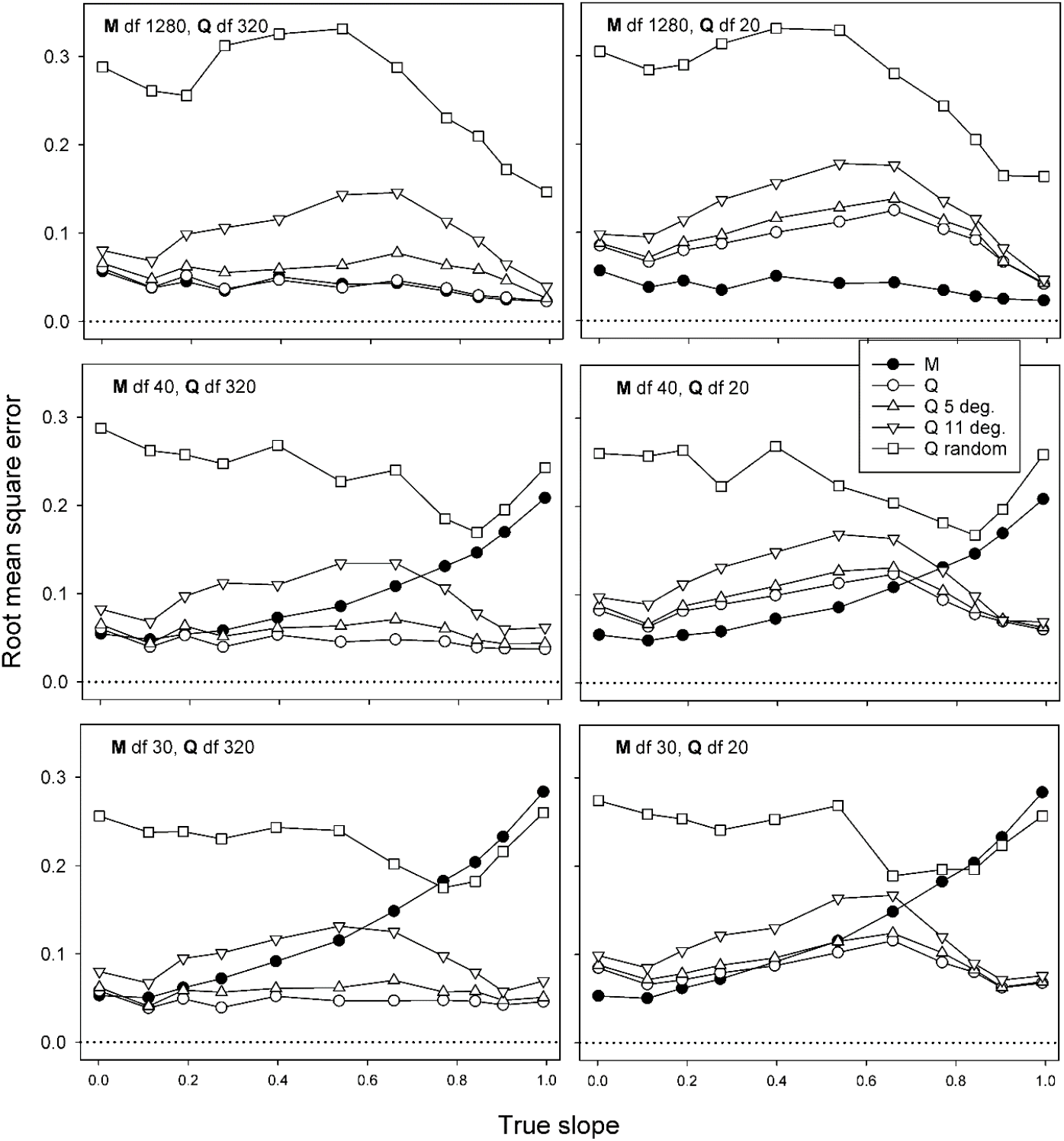
Geodesic distances between **M**_*i*_ and **Q**_*i*_ under various combinations of sampling error. Dashed reference lines show the range of distances among the observed matrices 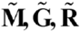 and 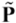 estimated from the data in H17. Legend as in Fig. 2.

### Neutral subset models

In H17, we suggested that a neutral-subset hypothesis could in principle explain the observed near-isometric relationships of variation. J&Z20 simulate a variant of the neutral subset model in which focal traits with a larger mutational target size are more constrained by deleterious pleiotropic effects on other traits than traits with smaller mutation target sizes. They show that such a model can generate scaling relationships with slopes that differ from 1. We now present an analytic variant of J&Z20’s model, and show that either hypometric or isometric scaling can result from variation in neutral mutational target size.

Assume that there is a probability *p_c_* that a new mutation has a deleterious pleiotropic effect, and that it will be eliminated if it has this effect. Then the conditional variance (Hansen and Houle 2008) in the trait would be *M_c_* = (1 – *p_c_*)*M*, where *M*is the unconditional mutational variance. To model a very high probability of having deleterious pleiotropic effects that is also increasing with number of loci, we use the function *p_c_* = Exp[−*dn*^−*k*^], where *n* is the number of loci affecting the trait, *d* is a small constant and *k* is a scaling exponent describing how fast the probability that a mutation is deleterious increases with number of loci. The small *d* yields the approximation

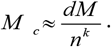

If all loci contribute the same mutational variance, 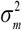, to the trait, 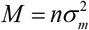, and we get 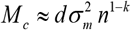. Under neutral evolution with mutational variance *M_c_*, the equilibrium genetic variance is *G* = *2NM_c_*, where *N* is the effective population size, and the equilibrium rate of evolution is *R* = *2M_c_* ((Lynch 1990)). Substituting 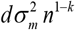 for *M_c_* and taking logarithms yields

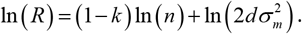

To obtain the scaling relationship with respect to *M* when *n* varies, we substitute 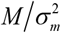 for *n* to get

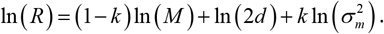

Applying this to a set of orthogonal traits, this suggests that the scaling exponent between variances in **R** (or **G**) and **M** would be 1 - *k*. Inspection of the model also shows that we get the same scaling with **M** if the variation is generated by differences in mutational rates or mutational effect sizes, 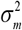, provided the probability of deleterious pleiotropy scale similarly (i.e. 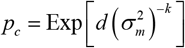). Hence, our observed near-isometric scaling is expected if the pleiotropic constraint is unrelated to the number of loci or to mutational effects. To get a scaling of, say, one half, we would need *k* = 1/2, which would imply that a doubling of the number of loci or mutational variance would increase the probability of being deleterious with a factor of 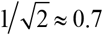. Note that the model predicts isometric scaling between **G** and **R**.

## Discussion

Jiang and Zhang (2020; abbreviated J&Z20) challenged some of the conclusions of our comparison of mutational variation, standing genetic variation and the rates of evolution of a suite of wing shape traits in the family Drosophilidae (Houle et al. 2017; abbreviated H17). Their first point is that the method that we used to choose traits on which to compare variation at these three levels, the H method, is inappropriate. J&Z20 show that results using the H method are insensitive to the true relationship between variation at the different levels. Our simulations, using a different method for varying the relationship between variance matrices, confirm J&Z20’s findings. The H method is clearly flawed, and should not be used.

Second, J&Z20 reanalyzed the relationship between mutational variation and the rate of evolution using the H17 data, and found that the scaling relationship was substantially lower, with a slope 0.54, rather than the slope of about 1 estimated by H17 using the flawed H method. In their analysis, J&Z20 used the mutational variance matrix, **M**, to choose the traits for comparisons. By doing so, they utilize the hypothesized predictor data to choose the traits for comparison, which we term the M method. This is known to introduce a downward bias to the estimated scaling relationship. The magnitude of the bias is a function of the discrepancy between the estimate of **M**, and the true variation, symbolized 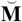. The bias arises because random discrepancies between **M** and 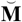 increase the range of the predictor variances that dependent variances are regressed on, decreasing the estimated slope.

For the data set utilized in H17, sampling variation is a minor source of discrepancy between **M** and 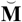. The relevant 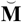 is the long-term average over the evolution of the family Drosophilidae, a period of at least 39.5 million years, and perhaps as many as 66 million years (Tamura et al. 2004; Grimaldi and Engel 2005; Wiegmann et al. 2011; Obbard et al. 2012; Russo et al. 2013). The **M** matrix used in H17 is the average of estimates from two different inbred lines with rather different estimates of **M** (Houle and Fierst 2013). These two genotypes were sampled from one natural population of one species, *Drosophila melanogaster*. Variation in **M** between inbred and outbred genotypes, among populations of *D. melanogaster*, and along the Drosophilid phylogeny all likely contribute to differences between **M** and 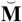.

As an alternative to the M method, we propose the Q method, which utilizes an independently estimated variance matrix, **Q**, for choosing traits along which to compare variation at different biological levels. Use of an independent matrix eliminates the bias inherent the M method, but can potentially affect the precision or bias of the estimates in other ways. This method is particularly attractive for the H17 data, as there are two other matrices that can be utilized as **Q**, the additive genetic variance matrix **G**, and the pooled phenotypic variance matrix **P**. We simulated the Q method for matrices with properties similar to the matrices estimated in H17. When the **Q** matrix is about as similar to the dependent and predictor matrices as those estimated in H17, the Q method has a modest positive bias at intermediate degrees of relationship between the predictor and dependent matrices, which decreases as dependent and predictor become more similar.

Clearly, whether the M or the Q method will perform better depends on the degree of difference between the estimated predictor matrix and its true value. We sought to determine which performed better for estimated matrices in H17 by seeking biological variances that match the scaling relationships estimated from the M and Q methods, the predictive power of the scaling relationships, and the overall similarity of all the estimated matrices. All these can only be explained when **M** and 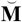 are substantially different. In such cases the expected bias and mean square-error in the estimated scaling relationship between **M** and **R** are substantially lower using the Q method than the M method. The use of **G** as **Q** leads to a scaling relationship between **M** and **R** of 1.01, while the use of **P** suggests a relationship of 1.13. Consequently, we conclude that the true scaling relationship between 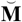 and **R** is closer to the original estimate of 1.02 given in H17, than to J&Z20s estimate of 0.54. This similarity of the H estimate to the Q estimates is not entirely fortuitous. One of the reasons that we accepted the flawed H method is that it yielded a very similar scaling relationship to the Q method using **G**, seeming to justify the H method.

We tailored our simulations of the M and Q methods to match the remarkable similarity in the overall pattern of eigenvalues of the matrices estimated in H17. Our inference that the Q method outperforms the M method is therefore not general. There are two aspects of the H17 matrices that suggest this conclusion. First, it appears that the true scaling relationship between all the H 17 matrices are near isometry where the Q method is almost unbiased. Second, our results indicate that there is a substantial bias to the M method for these data, suggesting that 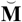 is rather different from the estimated **M** matrix. When the M method is applied to the relationship between **G** or **P** and **R**, the bias is substantially less, although still in the range in which the Q method is expected to perform better. These results need not hold for all sets of matrices. The relative performance of different methods of choosing a basis for matrix comparison will need to be reevaluated for new data sets.

The Q method is only applicable when there is a third matrix from which to estimate the basis. Furthermore, our simulations suggest that the Q method will only give reasonably accurate estimates when the **Q** matrix is similar to the predictor and dependent matrices. A potential alternative would be to fit a scaling model to entire matrices, rather than just to the trait variances as

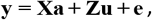

where **y** is the stacked vector of trait measurements, **X** and **Z** are fixed and random design matrices, **a** is the vector of trait grand means, **u** is the random effect of species deviations from the trait grand means, and **e** is a vector of residuals. To generate a single scaling coefficient, one would constrain the variation in **u** to be a matrix power function of the complete predictor variance matrix, for example **M**

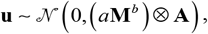

where *a* is a scalar, *b* is the exponent of the matrix power function, ⊗ is the Kronecker product, and **A** is the phylogenetic relationship matrix. To obtain the phylogenetic mixed model results in H17 and this paper, we fit a power function between predictor and dependent matrices when both matrices are constrained to be diagonal. As such, this required a choice of basis for the joint diagonalization, introducing the same biases discussed in this paper. The matrix-power approach would enable fitting scaling coefficients based on an a priori set of basis vectors not influenced by sampling of the estimates, bypassing the possibility of bias due to sampling. The logic of our mixed model approach in H17 is similar to the proposed matrix power approach, suggesting that it may be numerically feasible.

To explain the low scaling relationship between **M** and **R** that they observe, J&Z20 propose a model under which the proportion of mutations with deleterious pleiotropic effects increases with the mutational variance in a trait, while variation in wing shape itself is neutral. The pleiotropic mutations will be eliminated by natural selection, leaving the neutral subset to cause among-species variation. J&Z20 simulate evolution under this model, and show that this model can explain both scaling relationships of evolutionary rate with mutation with slopes less than 1, as their M estimates suggest, and the low rate of evolution shown in H17, as long as the proportion of neutral, non-pleiotropic mutations is small.

We have derived an analytical version of J&Z20’s model that confirms their simulation results. By varying how the proportion of neutral mutations varies with mutational variance, this model can explain any scaling relationship between **M** and **R**. An isometric scaling relationship, such as we observed is a special case where the proportion of mutational variation that is neutral is the same regardless of the amount of mutation variation in a particular direction in phenotype space.

This scenario of equal proportions of neutral variation in all phenotypic dimensions can be justified by a model of organ-specific effects. Assume, as in the J&Z20 model, that phenotypic variation in wing shape is effectively neutral, while variants with pleiotropic effects on other aspects of the phenotype are deleterious. Now further assume that only cis-regulatory variants with wing-specific effects can be neutral. Conversely, all coding variants have deleterious effects on non-wing phenotypes, as do regulatory variants with effects on both wings and other phenotypes. If the proportion of sites capable of affecting wing phenotypes that arise from wing-specific cis-regulatory machinery is approximately the same for all wing genes, then we can explain equal reduction in the effective mutational variation in all wing-shape traits.

J&Z20’s final point concerns the plausibility of various models that can potentially explain the results in H17. Plausibility is of course in the eye of the beholder. All we will add to this discussion is why we regard some models as more plausible than others. J&Z20 read H17 as suggesting that the neutral subset model was our favored explanation for the tight and near iso-metric scaling relationships we observe between variation at various biological levels. In fact, we termed this model implausible, although we did not explicitly state the basis for this opinion. We called it implausible due to the assumption that all wing shape variation is effectively neutral. For example, there is a clear signal of static and evolutionary allometry for wing shape in the Drosophilidae (Houle et al. 2003; Bolstad et al. 2015), and some of that allometry, such as the movement of veins to span an increasing proportion of the wing blade with increasing size make functional sense. Furthermore, the complex deformation of the wing during flight, and the degree of wing damage suffered by flies in nature suggest additional, if not yet experimentally verified, avenues for natural selection to directly influence wing shape. On the other hand, it seems unlikely that all aspects of wing shape are optimized by natural selection.

J&Z20 have done a service in pointing that our method of determining directions in which to compare variation is flawed, and should not be used. However, reanalyses based on the Q method we propose here confirms the findings in H17. Biologically plausible models of evolutionary rate, such as random walks of trait optima, do not fit the data. On the other hand, the models that do fit the data, the neutral subset models discussed above, or the vaguer hypothesis that mutation is reshaped to match the historical strength of selection, seem implausible. If our intuition that both selection and drift play a role in the diversification of the Drosophila wing shape, how can it be that the evolutionary rates that are any smooth function of mutational variation? This is the paradox of variation in the Drosophila wing.

## Acknowledgements

We thank Daohan Jiang and Jianzhi Zhang for sharing their comment with us prior to publication, the Centre for Advanced Study, Oslo for hosting DH and TH during the preparation of this manuscript, and members of the Evolvability Project at the Centre for feedback. During preparation of the manuscript, DH was supported by U.S. National Science Foundation (www.nsf.gov) Division of Environmental Biology grant 1556774 to DH and GHB by Norwegian Research Council project 287214 to Christophe Pélabon.

---

**Table S1.**
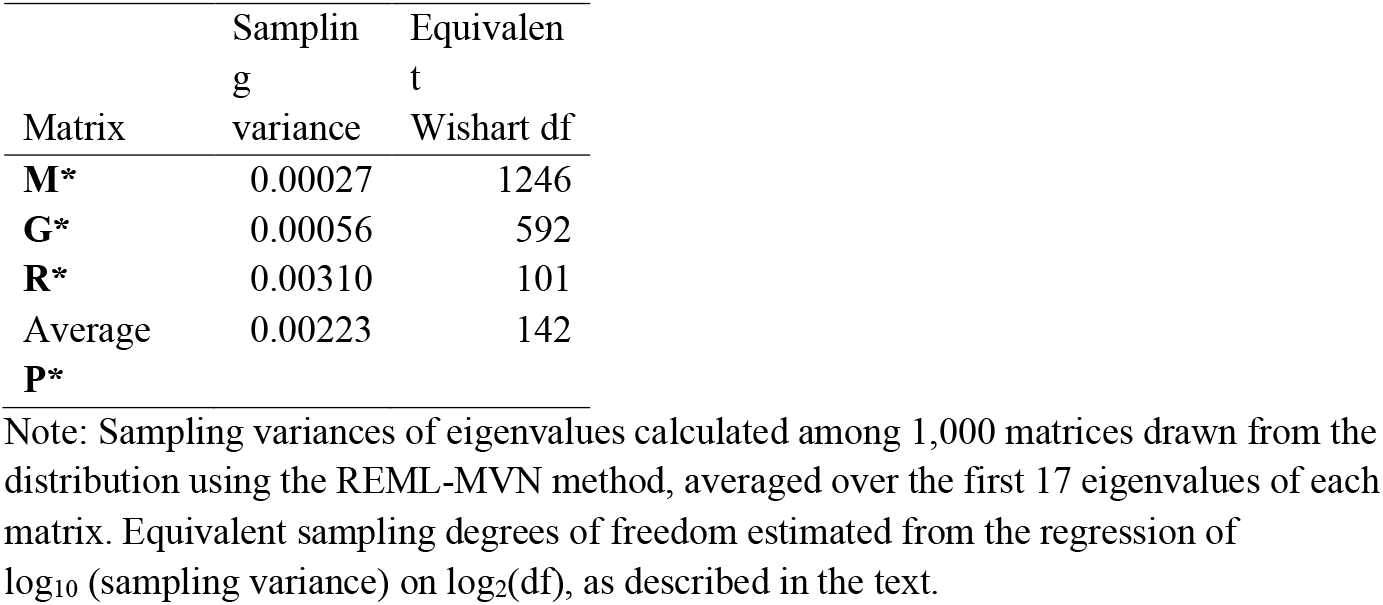
Sampling variances of log_10_(eigenvalues) in the standardized variance matrices estimated from the data in H17.

**Figure S1.**
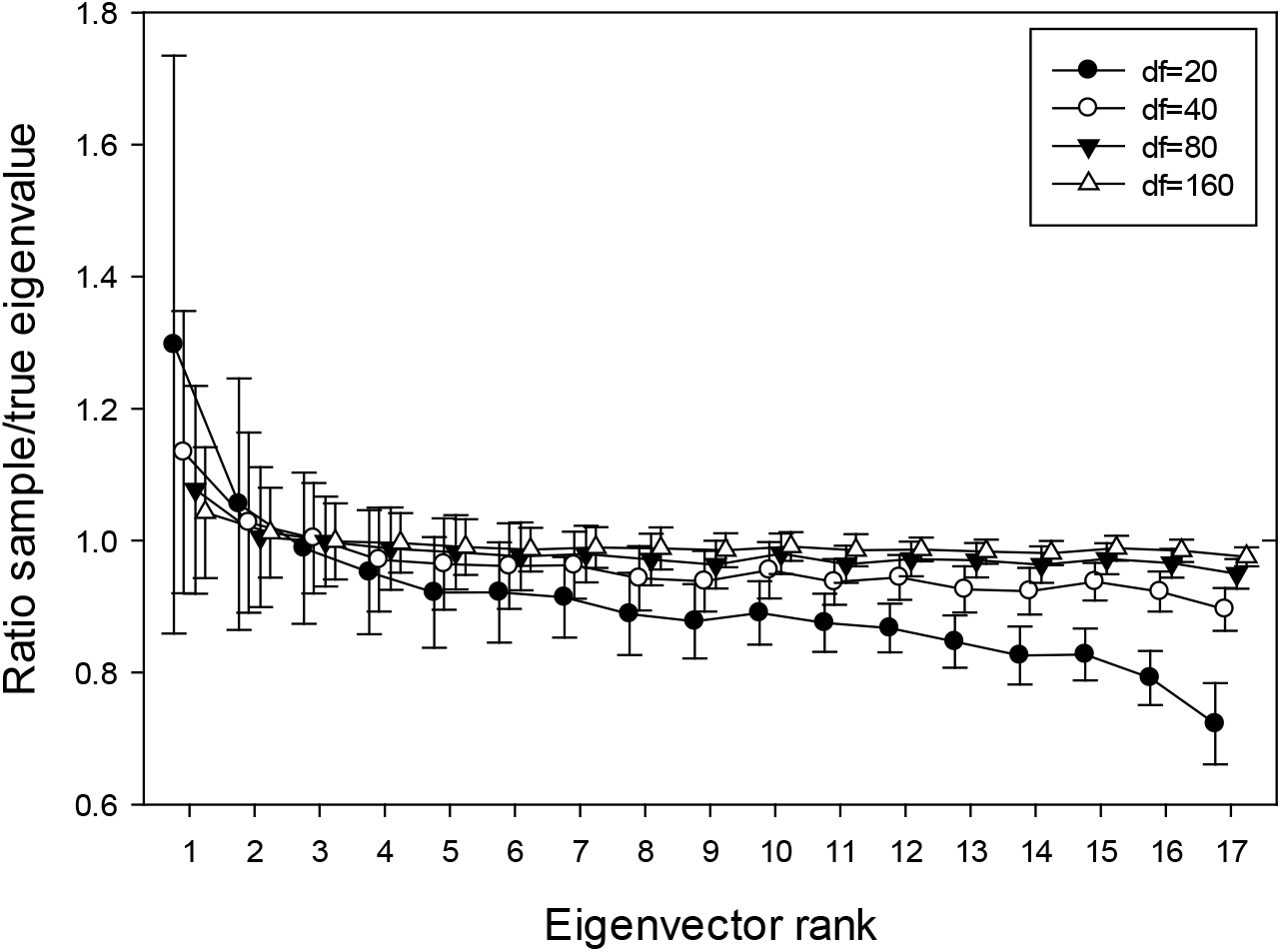
Ratio of sample to parameter eigenvalues ± standard deviation as a function of eigenvector rank and sampling variance. Parameter matrices were generated with a decline in log_10_(eigenvalues) with rank at the rate of *b*=-0.154 with sampling variance *v*=0.007 around that expectation. Results are the average of results from twenty parameter matrices, each sampled from a Wishart distribution 10 times with the df shown.

**Figure S2.**
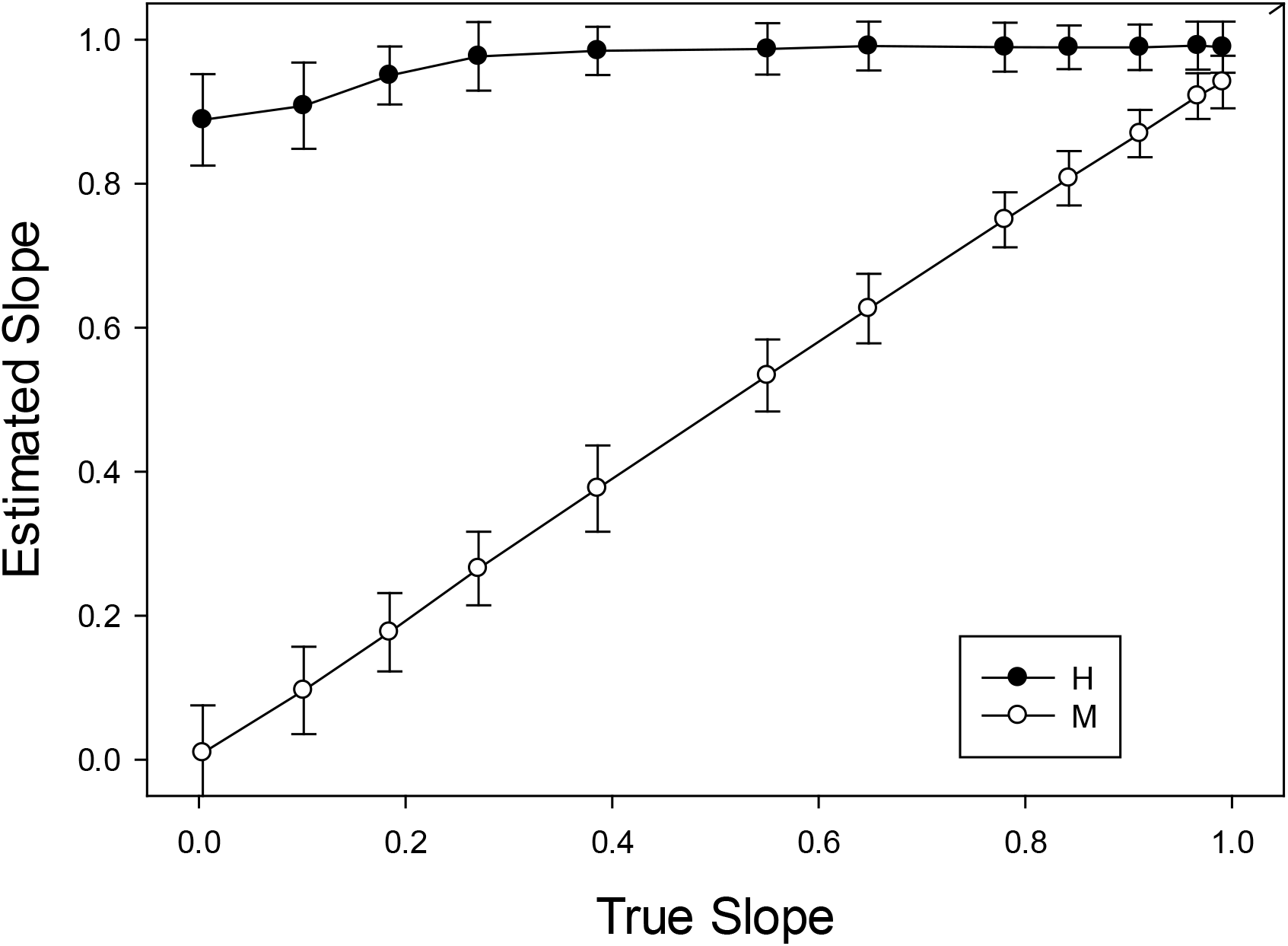
Estimated slope as a function of true slope for simulated matrices constrained to be similar to those in H17. Dashed line is the 1:1 line. For these results the sampling df=160 for both **M** and **R**.

**Figure S3.**
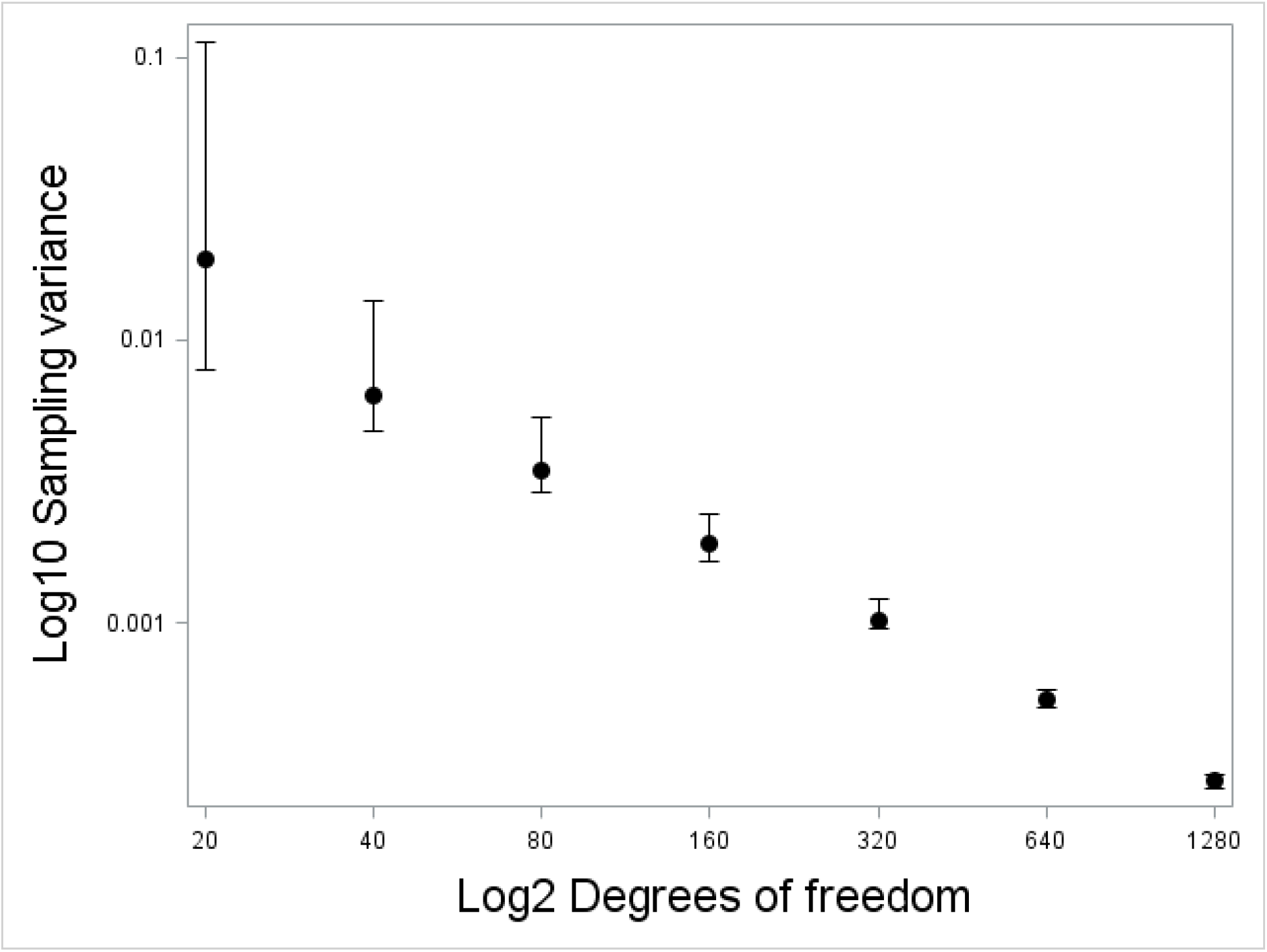
Distribution of the ratio of sample variance of eigenvalues to the variance in the parameter matrix as a function of the degrees of freedom during sampling. The ratio of 1.0 indicates no bias.

**Figure S4.**
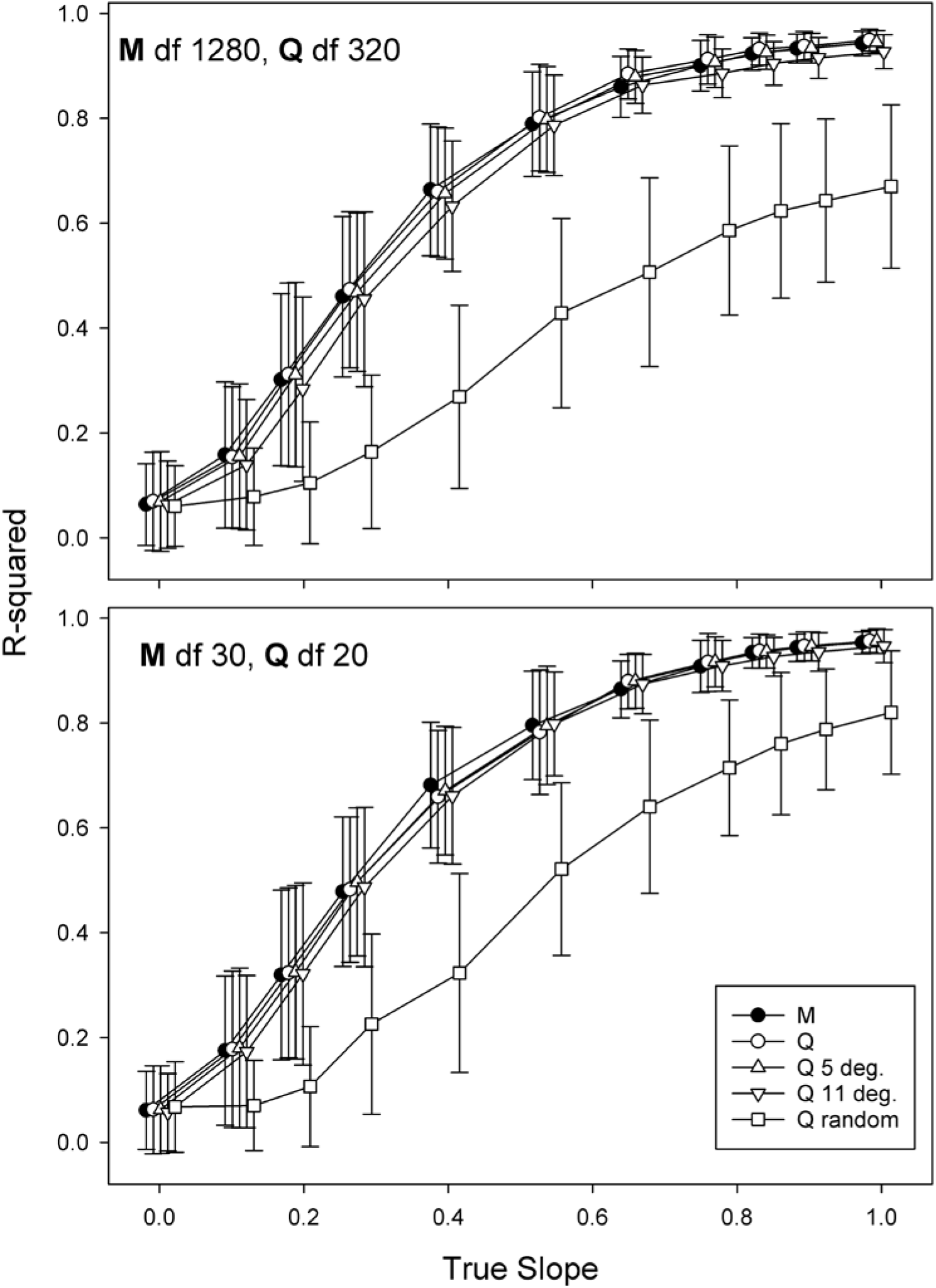
*R*^2^ of scaling relationships between **R**_*i*_ and **M**_*i*_ as a function of the true relationship when the M and Q methods are used to define a basis. True slopes are constant across basis assignment methods, symbols are offset to improve visibility. M = M method; Q = Q method when 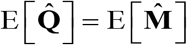; Q 5 deg. = as in Q, but 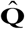 subjected to random rotations by 4.5 degrees; Q 11 deg. = as in Q, but 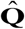 subjected to random rotations by 10.8 degrees; Q random = orientation of 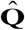 random with respect to 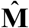.

